# Trapped DNA fragments in marine sponge specimens unveil north Atlantic deep-sea fish diversity

**DOI:** 10.1101/2023.03.29.534740

**Authors:** Erika F. Neave, Wang Cai, Maria Belén Arias, Lynsey R. Harper, Ana Riesgo, Stefano Mariani

## Abstract

Sponges pump water to filter feed and for diffusive oxygen uptake. In doing so, trace DNA fragments from a multitude of organisms living around them are trapped in their tissues. Here we show that the environmental DNA retrieved from archived marine sponge specimens can reconstruct the fish communities at the place of sampling and discriminate North Atlantic assemblages according to biogeographic region (from Western Greenland to Svalbard), depth habitat (80-1600m), and even the level of protection in place. Given the cost associated with ocean biodiversity surveys, we argue that targeted and opportunistic sponge samples – as well as the specimens already stored in museums and other research collections – represent an invaluable trove of biodiversity information that can significantly extend the reach of ocean monitoring.

## 1. Introduction

The worrying and widespread trend of ocean biodiversity loss that typifies the Anthropocene calls for increasingly powerful and accurate approaches to expose the nuances of this loss, understand its main drivers, and inform mitigation strategies. One such recent scientific advance has been ‘environmental DNA’ (eDNA) analysis, an approach by which collecting DNA fragments shed by organisms in their habitat, allows researchers to generate biodiversity data at unprecedented scales ^1^ and granularity ^2^, redefining the way we observe and understand ocean life.

Biological research collections are critical for eDNA analyses. Apart from expanding DNA taxonomic reference databases from tissues ^3^, they also provide untapped genomic insights that have become more accessible with the advancement of molecular techniques ^4^. Metabarcoding in particular allows for ecological insights, such as detecting multi-decadal community shifts from eDNA in ethanol-preserved ichthyoplankton samples ^5^, or tracking micro-evolutionary changes in the gut microbiome of 100-year-old fish specimens ^6^. These are prime applications of the extended specimen concept ^7^, that is, a novel, comprehensive approach to biodiversity collections that extends beyond the mere physical object to potentially limitless further uses that become possible owing to new efforts, such as digitization, and new attitudes towards phenotypic description ^8, 9^.

Filter-feeding marine sponges (phylum: Porifera) were recently found to act as natural eDNA samplers, able to retain eDNA fragments reflective of their surrounding communities ^10^. Sponges are ideal extended specimens, in that exploring beyond the host DNA provides an understanding of the environment from which the sponge was collected. Experimental studies subsequently found that sponge species differ in their ability to retain eDNA, with some species likely to trap DNA for longer intervals than what is usually observed in water samples ^11,12^. Given the urgent need to measure trajectories of biodiversity changes, we explored whether this sponge natural sampler approach could characterise fish assemblages across the North Atlantic, by leveraging sponge specimens previously collected for other scientific purposes from vulnerable and underexplored deep-sea habitats.

## 2. Results

We detected natural sampler DNA (nsDNA) from three sponge species (*Geodia barretti, Geodia hentscheli*, and *Phakellia ventilabrum*) (N = 54, retained from 64 samples sequenced – see Methods) across varied benthic habitats in the North Atlantic (Figure 1). The specimens were between 3-10 years old, spanning the continental shelf down to the bathyal slope (∼80-1900 m), and cover large biogeographic regions such as the Northeast Atlantic, North American Boreal, and Norwegian-Arctic Seas (Figure 1B, 1C, 1D) ^13^ (Supplemental Table 1). We amplified a fish-specific 12S mitochondrial rRNA marker (tele02) ^14^ from the previously extracted total DNA of the sponge specimens, and sequenced the targeted amplicons on an Illumina iSeq 100, resulting in 5,269,740 raw reads. After quality filtering (see Methods), we retained 4,565,067 reads for downstream analyses (Supplemental Table 2), resulting with a median of 12,992 reads per sample (N = 74) (Supplemental Figure 1), including controls (N = 10) and samples that were later removed (N = 10) for having low reads (mostly *G. hentscheli*).

**Figure 1.**
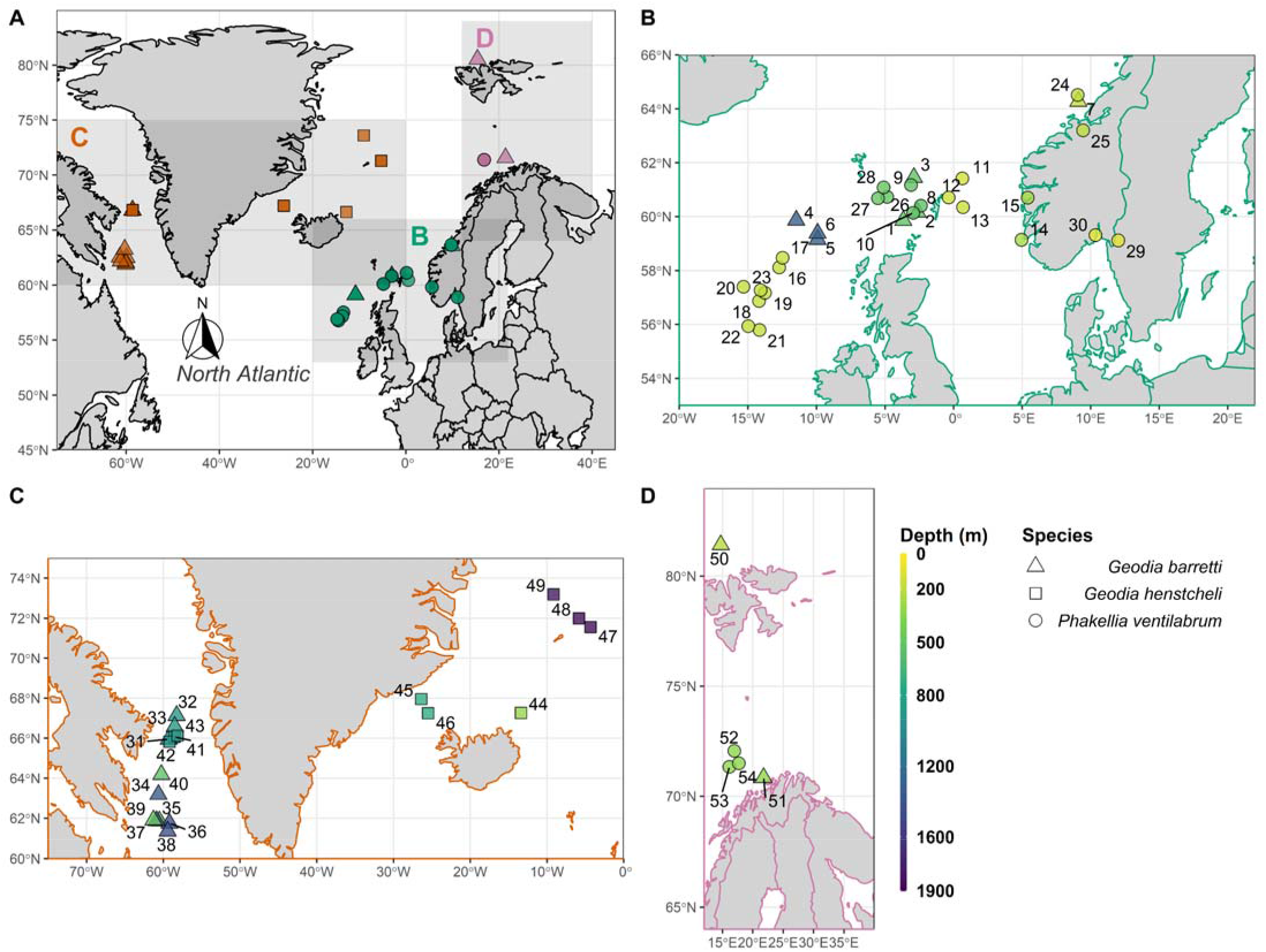
Maps showing locations of sponge specimen retrieval. Depth is indicated by the colour bar and sponge species is indicated by the shape of the points. **A** Panel showing the North Atlantic study area. **B** Panel showing the Northeast Atlantic region. **C** Panel showing the North American Boreal Atlantic region. **D** Panel showing the Norwegian-Arctic Seas Atlantic region. Sponge specimens in B, C and D are jittered for visibility and labelled 1-30 (Northeast Atlantic), 31-49 (North American Boreal) and 50-54 (Norwegian-Arctic Seas).

### (a) Vertebrate Biodiversity

The sponges yielded 142 eukaryote MOTUs, resulting in 125 non-human, contaminant free, marine MOTUs, which could be identified confidently to at least the taxonomic rank of class. Among these we detected 119 fish MOTUs of which 65 were identified to species level at ≥99% identity, excluding contaminants (Supplemental Table 2, 3). The following species were removed from downstream analysis: our positive control (the tropical freshwater catfish *Pangasianodon hypophthalmus*), two species (ie. *Amphiprion ocellaris, Pomacanthus imperator*) from a different project processed at a similar time ^12^, and one Indo-Pacific fish heavily traded as seafood (*Nemipterus zysron*). The fish MOTUs, spread over the classes Actinopterygii and Chondrichthyes, comprised 28 orders, 54 families and 94 genera. A sand sea star (*Astropecten irregularis*) common in deep sea benthos was also detected, while sponge DNA was never detected and likely not amplified, due to their phylogenetic distance from vertebrates. We also removed domestic animals (e.g. *Sus scrofa, Bos taurus*) and terrestrial mammals such as caribou (*Rangifer tarandus*), native to the Northern Hemisphere, whose putatively leached DNA was found in a *G. barretti* specimen from the Davis straight, west of Greenland. After these removals, we detected five ‘bonus’ non-fish vertebrate species, including three marine mammals (harbour porpoise (*Phocoena phocoena*), Atlantic white-sided dolphin (*Lagenorhynchus acutus*), and Bryde’s whale (*Balaenoptera brydei*) detected in both the west and east North Atlantic) as well as two seabirds (pelagic cormorant (*Phalacrocorax pelagicus*) and glaucous gull (*Larus hyperboreus*)).

### (B) Biogeography and depth-associated Fish assemblages

Fish communities significantly differed between biogeographic regions of the North Atlantic (R^2^ = 0.16, p < 0.001, Figure 2A, Supplemental Table 4). Beta diversity was examined through Non-metric Multi-Dimensional Scaling (NMDS) of a Jaccard dissimilarity matrix of teleosts and elasmobranchs detected across sponge samples, comprising of only MOTUs identified to the species level (though the same pattern resulted when including genus level detections, Supplemental Figure 2), and by permutational multivariate analysis of variance (PERMANOVA) testing. Sponge samples appeared broadly grouped into the biogeographic regions previously determined from global distribution data of marine taxa ^13^, emphasizing the effectiveness of sponge nsDNA to capably distinguish between marine realms (Figure 2A). Pairwise comparisons of beta-diversity revealed that all regions significantly differed, with the North American Boreal region showing greater divergence from both the Northeast Atlantic (R^2^ = 0.14, p < 0.001) and the Norwegian-Arctic Seas (R^2^ = 0.13, p < 0.001), compared to the divergence observed between the regions located in the eastern North Atlantic (R^2^ = 0.06, p = 0.025) (Supplemental Table 4).

**Figure 2.**
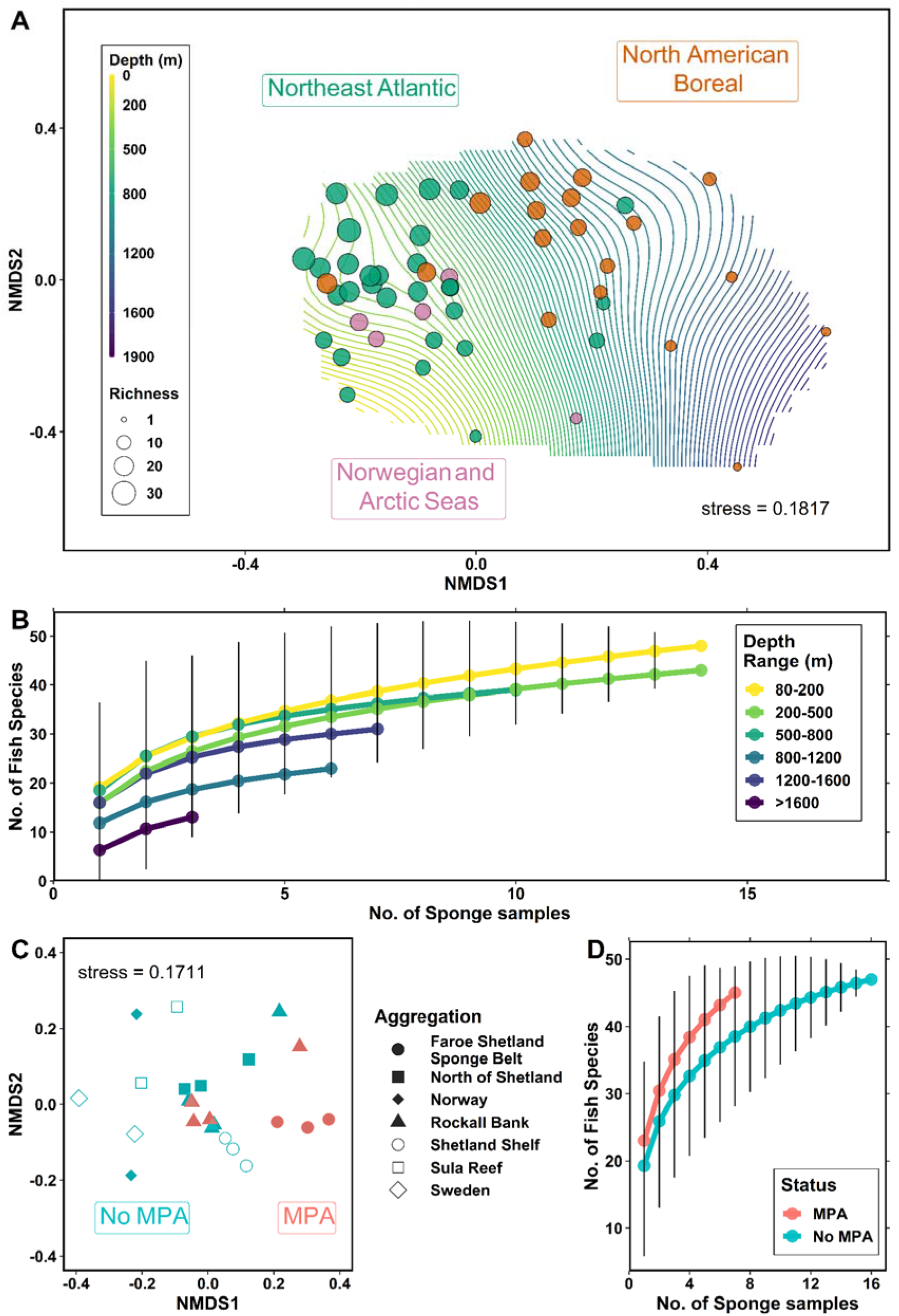
Plots conveying alpha and beta diversity from species-level teleost and elasmobranch detections. **A** Non-metric Multi-Dimensional Scaling (NMDS) plot of a Jaccard dissimilarity species matrix, where points are coloured by North Atlantic region and size indicates species richness. Depth is plotted as a surface, where each line denotes a 20 m interval. **B** Fish species accumulation curve, grouped by depth range. **C** NMDS plot of a Jaccard dissimilarity species matrix of *P. ventilabrum* samples from the Northeast Atlantic region. Points are coloured by MPA status and shapes represent different sponge aggregations. **D** Fish species accumulation curve of the *P. ventilabrum* samples from the Northeast Atlantic region, grouped by MPA status.

Latitude, depth and sampling year were all significant correlates of fish beta-diversity. Depth had the strongest correlation (R^2^ = 0.58, p < 0.001) followed by latitude (R^2^ = 0.35, p < 0.001) and year (R^2^ = 0.17, p = 0.018) (Supplemental Table 4). We attribute the weaker correlation with sampling year to be a by-product of the different regions being sampled in separate years. Depth was plotted as a smooth surface over the NMDS ordination plane (Figure 2A), particularly highlighting how the composition of the Northeast Atlantic sites correspond with shallower continental shelf depths, while the North American Boreal samples follow the gradient of the slope into bathypelagic depths. Species richness approached saturation among all depth ranges from which sponges were sampled, but more robustly in shallower groups (80-200 m) that had a greater sample size. Fish species richness progressively decreased with depth, except between 1200-1400 m depth, which had a higher richness than the 800-1200 m samples, but also had a greater sample size (Figure 2B).

To further test the extent to which sponge nsDNA data could be used to distinguish between more fine-scale fish assemblages, the *P. ventilabrum* samples from the Northeast Atlantic were analysed as a subset (N = 23) to compare similar habitats and to control for any possible bias introduced by using different sponge species. We observed variance across samples collected in areas with differing levels of marine protection. Species richness appeared to be higher in marine protected area (MPA) sites, and communities detected in MPAs significantly differed from those outside MPAs (R^2^ = 0.09, p = 0.026) (Figure 2C, 2D). The same subset of sponges was also tested for significant differences in teleost and elasmobranch beta-diversity between various *P. ventilabrum* aggregations (Figure 2C); however, none of the pairwise comparisons among aggregations were significant after correcting the p-values for multiple testing (Supplemental Table 5). This was likely due to low replication within each of the several locations (e.g., Sula reef, Shetland Shelf) being compared.

### (c) Fish detections and indicator species analysis

Greenland halibut (*Reinhardtius hippoglossoides*), beaked redfish (*Sebastes mentella*), and megrim (*Lepidorhombus whiffiagonis*) were detected in almost all 54 samples (i.e., 52, 51, and 50 samples, respectively) (Figure 3, Supplemental Table 3). Other frequently detected species included Atlantic mackerel (*Scomber scombrus*), greater argentine (*Argentina silus*) and poor cod (*Trisopterus minutus*) (i.e., 48, 46, and 44 samples, respectively), all of which are known to be abundant organisms in pelagic and demersal habitats of the North Atlantic.

**Figure 3.**
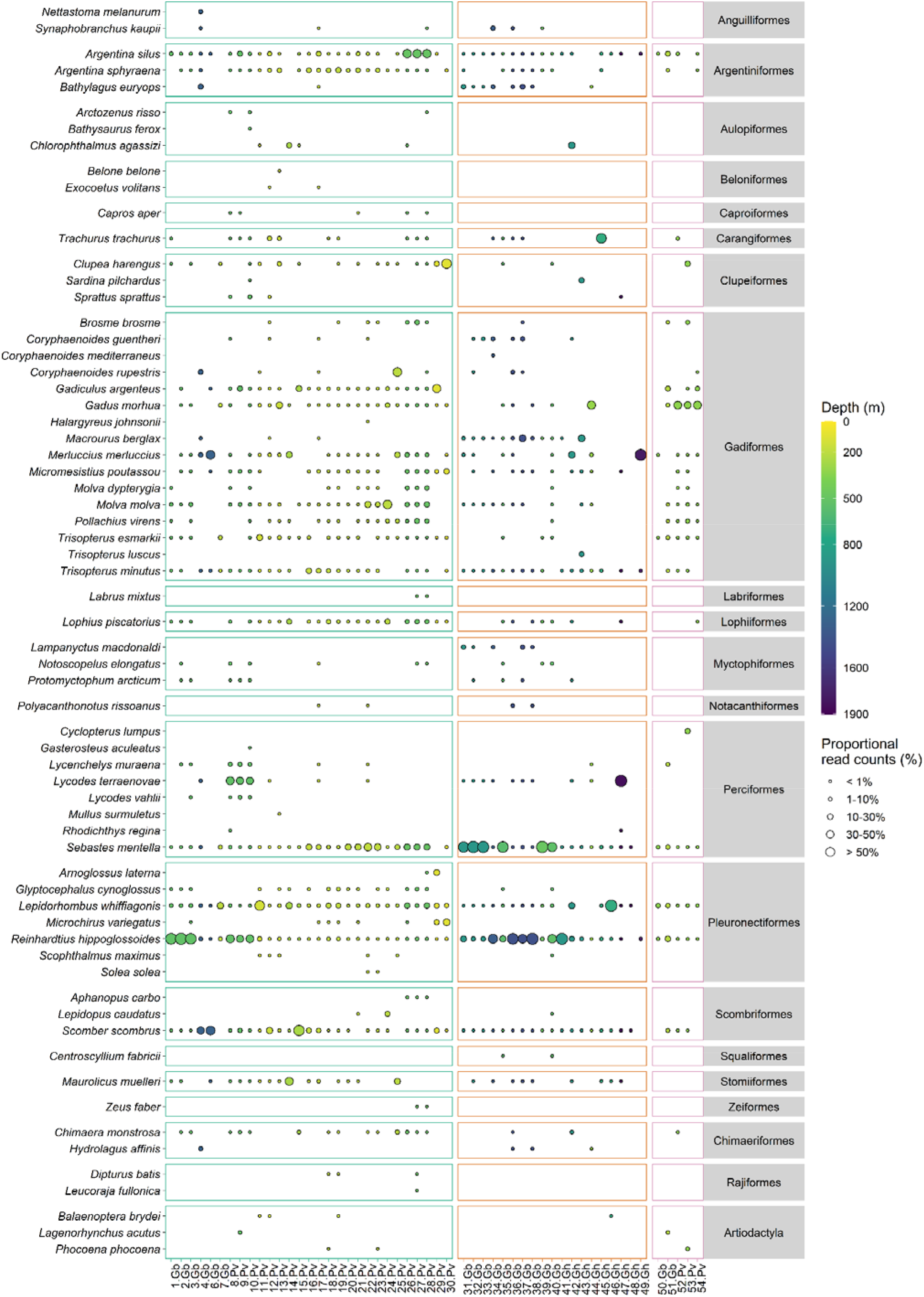
Bubble plot showing teleost, elasmobranch and mammal species detected, where the size of the bubble indicates the proportional read counts of that species represented in a sample. Samples are listed at the bottom, where the number refers to the labels in Figure 1 and the abbreviations refer to the sponge species (Gb = *Geodia barretti*, Gh = *Geodia hentscheli*, and Pv = *Phakellia ventilabrum*). The bubbles are coloured by the depth at which the sponge specimen was sampled. The panels separate the sponge specimens by the three biogeographic regions and are coloured accordingly: Green = Northeast Atlantic, Orange = North American Boreal, and Pink = Norwegian-Arctic Seas (Same colour scheme as Figure 2A).

While the 12S marker was designed to pick up teleost fish, six cartilaginous fish (class: Chondrichthyes) were also detected. Three chimaeras, the closest living relatives to sharks and rays, were detected, including the rabbit fish (*Chimaera monstrosa*) which was detected in 17 samples.

Two elasmobranchs were from the family Rajidae: the shagreen ray (*Leucoraja fullonica*) which is IUCN red-listed as vulnerable and the blue skate (*Dipturus batis*) which is critically endangered, were both detected in the Northeast Atlantic (Figure 3).

Indicator value species analysis conducted across biogeographic regions and depth ranges (Figure 4A, 4B) detected eight species as biogeographic indicators, and 16 species as depth layer indicators, with seven species identified as indicators for both region and depth (Supplemental Table 5).

**Figure 4.**
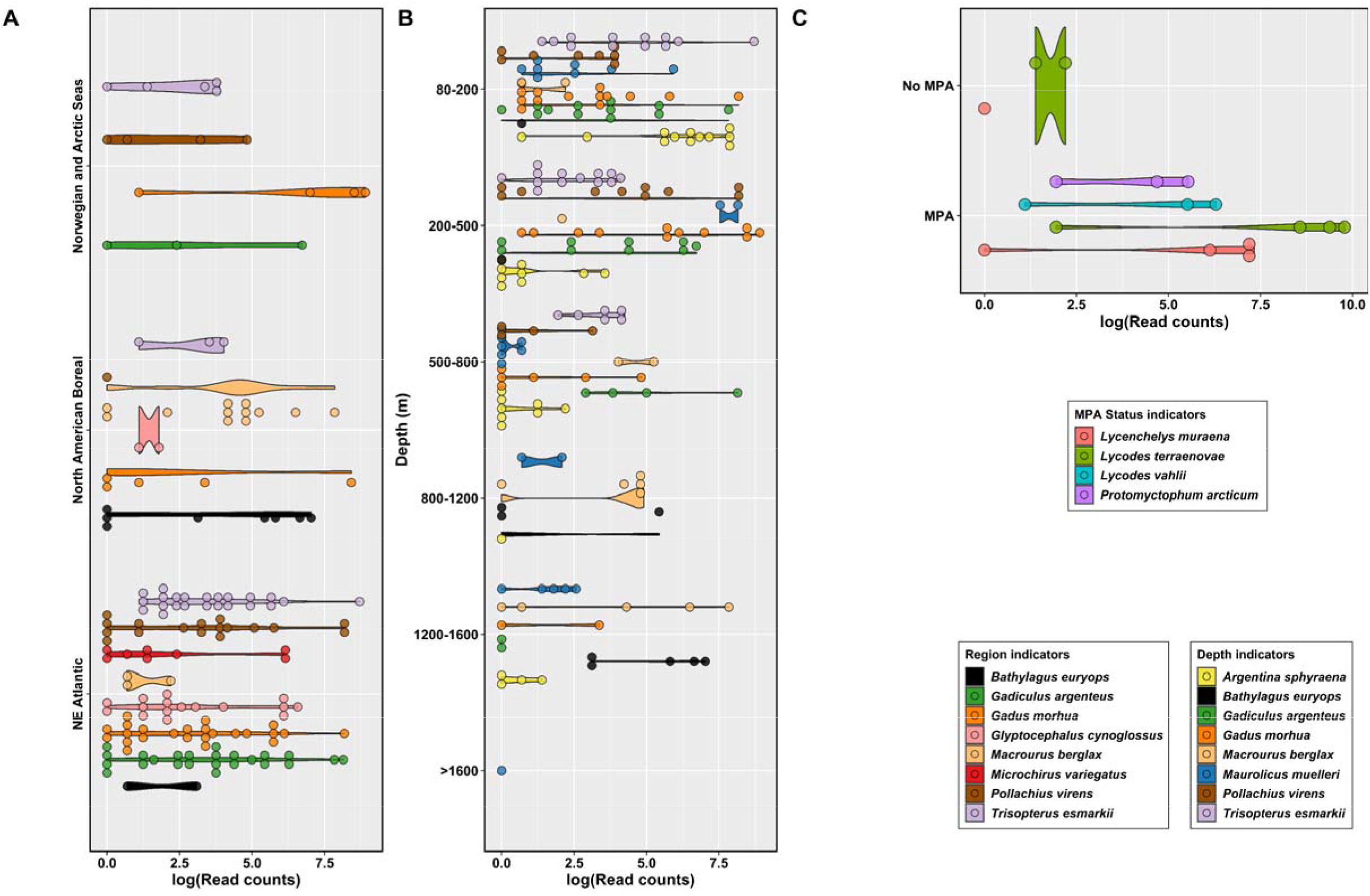
Violin dot plots of log-transformed read counts, highlighting identified indicator species. A Violin dot plot of indicator species associated with biogeographic regions. B Violin dot plot of the top eight indicator species associated with depth. C Violin dot plot of indicator species associated with MPA status.

Indicator values (A, B, stat) were calculated using presence-absence data to conservatively interpret detections. “A” is the estimate probability that samples are associated to a region or depth layer if the indicator species has been detected in the sample (i.e., specificity or predictive value). “B” is the estimate probability of detecting the indicator species in a region or depth layer (i.e., sensitivity). “Stat” is the indicator value index which suggests the strength of the indicator species association and encompasses both “A” and “B” values.

Many species of commercial value had strong significant associations for both region and depth range. Norway pout (*Trisopterus esmarkii*) was positively associated with the Northeast Atlantic and Norwegian-Arctic Seas (stat = 0.857, p < 0.0001) (Figure 4A) and had a strong association with depths ranging from 80-800 m (stat = 0.903, p < 0.0001), such that there was high specificity (A = 1) or likelihood that a Norway pout detection occurred in habitats shallower than 800 m depth (Figure 4B, Supplemental Table 6). Pollock (*Pollachius virens*) shared the same region and depth associations as Norway pout, although to a lesser strength. Atlantic cod (*Gadus morhua*) also showed clear associations with the eastern Atlantic between 80-800 m (Figure 4A, 4B). Roughhead grenadier (*Macrourus berglax*) and blacksmelt (*Bathylagus euroyops*) had a strong association with the North American Boreal with grenadier having a higher likelihood of detection (B = 0.738) than blacksmelt (B = 0.603). Both species were associated to depths between 800 and 1600 m (Figure 4B).

The indicator species analysis was repeated with the *P. ventilabrum* subset (N = 23) of the Northeast Atlantic data to identify indicator species of MPA sites. Four species were significant indicators of MPA sites (Figure 4C). These species included the moray wolf eel (*Lycenchelys muraena*), Atlantic eelpout (*Lycodes terraenovae*), Arctic telescope (*Protomyctophum arcticum*) and Vahl’s eelpout (*Lycodes vahlii*), all of which had high specificity (A = 0.999, 0.999, 1.0, and 1.0, respectively) to MPAs. The moray wolf eel and the Atlantic eelpout both shared the highest association with MPAs (stat(s) = 0.756, p < 0.05) (Supplemental Table 6).

## 3. Discussion

The retrieval of fish sequences from sponge specimens previously collected for other monitoring purposes provides perhaps the most attractive demonstration to date of the role of sponges as practical, cost-effective, universal natural DNA samplers for aquatic biodiversity studies. We confidently detected at least 65 teleost and elasmobranch species that could be used to distinguish fish assemblages and identify indicator species associated with depth and biogeographic regions within the North Atlantic.

Congruent with what we know about sponge nsDNA *ex situ* ^12^, some sponge species appeared to perform better than others. The original experimental design considered 93 sponge specimens; however, only 64 of them were selected for sequencing because they showed amplification of the desired target DNA region (i.e., bands on agarose gels). After bioinformatic quality control, DNA information from 54 individual sponges was retained. Of the 34 *G. hentscheli* samples attempted, only 17 were sequenced and nine were kept after rarefaction. Sample loss occurred, although to a lesser degree, also for *G. barretti* (i.e. 33 attempted, 21 sequenced, 19 kept). *P. ventilabrum* resulted instead in a 100% success rate (N = 26), followed by *G. barretti* (58%) and *G. hentscheli* (26%).

Curiously, *P. ventilabrum* likely has higher pumping rates and lower microbial abundance than the *Geodia* species ^15,16^. It is possible that higher microbial abundance could contribute to increased rates of eDNA decay within sponges due to decomposition by bacteria ^15^ and less need to derive energy from the uptake of dissolved organic carbon ^17^. Given these observed coincidences, relationships between sponge physiology and nsDNA efficacy would be an exciting area for further investigation.

High DNA sampling efficiency in some sponge species (i.e. *P. ventilabrum*) is an obvious advantage for biomonitoring, yet the percent success rate of the tetractinellid (*Geodia*) sponges was comparable to or even better than other organisms that have been tested as natural DNA samplers. For example, various leech species have been used to detect prey DNA, with vertebrate detection rates ranging from 9% to 80% of attempted specimens ^18^. Similarly, when gut contents of the European brown shrimp (*Crangon crangon*), a generalist scavenger, were analysed with DNA metabarcoding to reconstruct estuarine fish assemblages ^19^, up to eight stomachs had to be pooled, per DNA extraction, to constitute a sufficient sample. Extraction pooling could represent an appropriate methodological solution for favourable and widespread sponge species with moderate amplification success, such as *G. hentscheli* (i.e. 26%).

The detected fish communities significantly differed between biogeographic regions of the North Atlantic (Figure 2A), and depth was identified as the most important variable in shaping beta-diversity (Supplemental Table 4). Several fish species seemed to be more associated with either the west or east North Atlantic. Thickback sole (*Microchirus virens*) was unique to the Northeast Atlantic; saithe (*Pollachius virens*) and Norway pout (*Trispoterus esmarkii*) were present in the east Atlantic far more than the North American Boreal. Seven species, most of commercial value, were identified as significant indicators of both region and depth. Fishes known to be deep-sea adapted were indeed significantly associated with greater depths, for instance, Rakery beaconlamp (*Lampanyctus macdonaldi*) from 800-1600 m and small-eyed rabbitfish (*Hydrolagus affinis*) from 1200-1600 m.

Moreover, the mesopelagic silvery lightfish (*Maurolicus muelleri*) was significantly associated with all sampled depth layers, except for 200-500 m, suggesting that the nsDNA signal detected their flexible migratory behaviour ^20^. Interestingly, Atlantic cod (*Gadus morhua*) was associated with the same region and depth range as the silvery pout (*Gadiculus argenteus*), which could be indicative of their known predator-prey relationship ^21^.

Fish assemblages under different MPA status were distinguishable within the subset of *P. ventilabrum* specimens from the Northeast Atlantic, and greater species richness was observed in specimens from MPAs (Figure 2C, 2D). Indicator species associated with MPAs were mostly benthic, such as the moray wolf eel (*Lycenchelys muraena*), which prey on crustaceans and other invertebrates that take refuge in sponge grounds ^22^. Atlantic eelpout (*Lycodes terraenovae*) and Vahl’s eelpout (*Lycodes vahlii*) were also indicators and known to eat sponge remains and cryptofaunal organisms such as brittle stars ^23^. *Lycodes* sp. have also been found to correlate positively with high sponge biomass ^24^. Differences between the sponge aggregations were strong (R^2^ = 0.52) though when pairwise comparisons were made, only one pair, the Faroe Shetland Sponge Belt and Rockall Bank, was identified as a potential driver of the difference. While the significant difference between MPA status was modest (R^2^ = 0.09), with adequate samples sizes and targeted rather than opportunistic sampling, sponge nsDNA shows promise for more fine-scale biodiversity surveying.

Environmental DNA analysis is an emerging tool for deep-sea biodiversity ^25–27^ and ecological studies ^28–30^, yet eDNA is less abundant in the deep-sea, such that larger volumes of water are needed to attain representative samples, and the manual labour required to filter those samples *in situ* can become a limitation ^31^. Furthermore, remote, deep-sea habitats are expensive to reach in the first place, so leveraging of natural samplers in this context represents a major boost for large scale ocean exploration and monitoring. For instance, the specimens in this study had previously been used to understand sponge phylogenetics and connectivity of deep-sea environments ^32,33^. The deep sea and high seas are subject to threats such as overfishing ^34^, deep-sea mining ^35^, climate change and pollution ^36^. Sponges are habitat-forming organisms ^37^ that provide shelter for cryptic animals, thereby also attracting larger more mobile predators ^38^, and as such playing a fundamental role in the structure and functioning of marine ecosystems. Now, the wealth of environmental, biological and molecular data that can be comprehensively obtained from sponges significantly expands their broader value in marine ecology and conservation.

## 4. Methods

### (a) Specimen Selection

Three sponge species: *Phakellia ventilabrum*, N = 26 (order Bubarida); *Geodia barretti*, N = 21 and *Geodia hentscheli*, N = 17 (order Tetractinellida) from various North Atlantic sponge grounds were selected for sequencing (N = 64 of which 54 were analysed for the study – see Statistical Analysis), all collected previously for the SponGES project (www.deepseasponges.org), which ran until 2020 (Supplemental Table 1). The sponges were stored in 100% EtOH which was replaced at least once to maintain a high percentage of EtOH, since the water retained by the sponges can significantly dilute the preservative. The sponge DNA had been extracted between 6 and 36 months after sampling using the Qiagen DNeasy Blood and Tissue Kit (Hilden, Germany), optimal for sponge nsDNA extraction ^39^ and were stored at the Natural History Museum at -80°C until being transported to - 20°C freezers at Liverpool John Moores University. These samples have now been deposited in the Natural History Museum collections under voucher numbers found in Supplemental Table 1.

### (b) Library Preparation and Sequencing

DNA extracts were diluted with molecular grade water to between 30-50 ng/μl. DNA was amplified using PCR with the Tele02 primers ^14^. The forward sequence Tele02-F (5’-AAACTCGTGCCAGCCACC- 3’) and the reverse sequence Tele02-R (3’-GGGTATCTAATCCCAGTTTG-5’), were used to target a 167 bp fragment of the mitochondrial 12S rRNA gene. PCRs were prepared to a total volume of 20 μl for each sample and included 10 μl of 2X MyFi Mix (Meridian Bioscience), 1 μl of each forward and reverse primer, 0.16 μl Bovine Serum Albumin (Thermo Fisher Scientific), 5.84 μl molecular grade water, and 2 μl of diluted DNA extract. The samples were amplified in triplicate across two libraries using the following conditions: 95°C for 10 min, followed by 35 cycles of 95°C for 30 s, 60°C for 45 s, 72°C for 30 s, and finishing at 72°C for 5 min followed by a 4°C hold. Negative controls (N = 5) and positive controls (N = 5), which were molecular grade water and a single fish species not present in the North Atlantic (iridescent catfish *Pangasionodon hypopthalmus*) respectively, underwent PCR alongside the samples. PCR triplicates were pooled and visualized on a 2% agarose gel (150 ml 1X TBE buffer with 3 g agarose powder) stained with 1.5 μl SYBRsafe dye. PCR products were individually purified using a double-size selection in 1:1 and 0.6:1 ratio of Mag-Bind® Total Pure NGS magnetic beads (Omega Bio-Tek) to PCR product. Products were visualised on an agarose gel again to assure purity (i.e., target length bands on agarose gels were visible with minimal to no other bands present). Purified PCR products were quantified using a Qubit dsDNA HS Assay kit (Invitrogen), and pooled equimolar into their corresponding libraries (i.e., pooled samples each contained unique 8-bp dual barcodes). Pooled libraries were imaged on a Tape Station 4200 (Agilent) to check the purity of the libraries. The libraries were then purified based on the Tape Station results, double-size selecting the target fragment using magnetic beads as explained before. A unique adapter sequence was ligated to each library using the NEXTFLEX® Rapid DNA-Seq Kit for Illumina (PerkinElmer) following the manufacturer protocol. After adapter ligation, the libraries were again imaged on the Tape Station and purified with magnetic beads, this time with a 0.8:1 ratio of beads to sample, as per the NEXTFLEX® Rapid DNA-Seq Kit instructions. The dual-indexed libraries were then quantified by qPCR using the NEBNext® Library Quant Kit for Illumina (New England Biolabs). The libraries were pooled at equimolar concentrations having a final molarity of 50 pM with a 10% PhiX spike-in. The libraries were sequenced at Liverpool John Moores University on an Illumina iSeq100 using iSeq i1 Reagent v2 (300 cycles).

### (c) Bioinformatics Pipeline

The sequences were quality controlled through the following series of steps using Python v2 within the OBITOOLS 1.2.11 ^40^ package. The raw sequences were trimmed to a length of 150 bp using the command ‘obicut’ to remove low-quality bases from the ends which were determined from the output of the ‘fastqc’ command. The trimmed reads were then merged using ‘illuminapairedend’, from which any paired-end alignments with low (<40) quality scores were removed. The remaining paired-end alignments were demultiplexed using ‘ngsfilter’, filtered by length (130 - 190 bp) and dereplicated using ‘obiuniq’. Chimeras were removed de novo using the programme VSEARCH version 2.4.3 ^41^. The remaining sequences were then clustered using the programme SWARM v2 ^42^ with ‘d-value’ = 3. Taxonomy was assigned using the Bayesian LCA-based taxonomic classification method (BLCA) ^43^. We first created a database using ‘ecoPCR’ from OBITOOLS with the Tele02 primers against the EMBL database (release version r143). This database was combined with a trained BLCA custom database containing fish species, specifically Teleosts and Elasmobranchs, (custom database file can be found here: https://github.com/eneave/Trapped-DNA-fragments-in-marine-sponges-Neave-et-al-2023). The workflow of BLCA was followed and can be found at: https://github.com/qunfengdong/BLCA. This resulted in taxonomic assignments where each level (i.e., family, genus) was associated with a percent probability of correct assignment. Analyses were carried out with taxonomies that had a ≥ 99% probability of correct assignment (i.e., species referenced in this study had a ≥99% identity at the species level and 100% identity at all higher levels of assignment).

### (d) Statistical Analysis

All downstream analyses were done using R version 4.1.3 ^44^. The MOTUs were decontaminated by removing the highest number of reads of a contaminant present in either the PCR positive control or PCR negative control from all samples (Supplemental Figure 3). Ten samples that had less than 100 reads were removed from the dataset based on a rarefaction curve suggesting species saturation after 100 reads (Supplemental Figure 1). Using the R package vegan v 2.5.7 ^45^, beta-diversity was examined through multi-dimensional scaling of a Jaccard dissimilarity matrix of teleosts and elasmobranchs detected from each sponge, comprising of only MOTUs identified to the species level. We tested the homogeneity among the group dispersions of biogeographic regions using the functions ‘betadisper’and ‘anova’, then tested for significant differences in beta-diversity between regions by permutational multivariate analysis of variance (PERMANOVA) using the function ‘adonis’. The same tests were repeated for the *P. ventilabrum* subset of the Northeast Atlantic. Pairwise comparisons of the biogeographic groups and population groups were performed, and p-values were corrected with the Benjamini-Hochberg method ^46^. Correlations of fish assemblages with latitude, sampling depth, and sampling year were tested for using the function ‘envfit’. All tests on beta-diversity were done on Jaccard dissimilarity matrices and underwent 1000 permutations. The ‘accumcomp’ function from the BiodiversityR package v 2.14.2.1 ^47^ was used to create species accumulation curves. Using the R package indicspecies v 1.7.12 ^48^, an indicator value species analysis and multilevel pattern analysis was done using the function ‘multipatt’ with IndVal.g method on the same Jaccard dissimilarity matrix of species for sampling depth ranges, biogeographic regions and MPA status in the Northeast Atlantic with *P. ventilabrum* samples. Tests underwent 10,000 permutations. All figures were generated using the R packages tidyverse v 1.3.1 and ggplot2 v 3.4.0 ^49,50^. All raw data and code can be found through the links in the data accessibility statement.

## Data accessibility

The raw data files can be accessed here: https://doi.org/10.5281/zenodo.7740858. The code used to process the data can be found in the following GitHub repository: https://github.com/eneave/Trapped-DNA-fragments-in-marine-sponges-Neave-et-al-2023.

## Author’s Contribution

Conceptualization, E.F.N., M.B.A., L.R.H., A.R. and S.M.; Investigation, E.F.N, M.B.A and L.R.H.; Formal Analysis, E.F.N. and W.C.; Visualization, E.F.N.; Writing – Original Draft, E.F.N. and S.M.; Writing – Review & Editing, E.F.N., W.C., M.B.A., L.R.H., A.R. and S.M.; Supervision and Funding Acquisition, A.R. and S.M.

## Competing Interests

We declare that we have no competing interests.

## Funding

The study was supported by Grant NE/T007028/1 (*SpongeDNA*) from the UK Natural Environment Research Council to S.M. and A.R. A.R. was also supported by grants RYC2018-024247-I and PID2019-105769GB-I00 from the Spanish Ministry of Science and Innovation, in the framework of both MCIN/AEI/10.13039/50110001103 and “FSE invierte en tu futuro”, and an intramural grant from CSIC (PIE-202030E006).

## Acknowledgements

We thank Ellen Kenchington, Jim Drewery, Vasiliki Koutsouveli, Hans Tore Rapp, Joana Xavier, Paco Cárdenas, Nathan J. Kennny, Kathrin Busch, and Karin Steffen, for facilitating access to the samples analysed here, in the framework of the SponGES project (grant ID: 679849). We are also indebted to Sergi Taboada, Alex Cranston, Alex Mitchell, Connie Whiting, and Karin Steffen, who performed DNA extractions. We thank Peter Shum for his assistance with the Illumina iSeq100 system.

## Notes

https://doi.org/10.5281/zenodo.7740858

https://github.com/eneave/Trapped-DNA-fragments-in-marine-sponges-Neave-et-al-2023

### Competing Interest Statement

The authors have declared no competing interest.

### Summary of Updates

The middle initial of Stefano Mariani's name has been removed because it is incorrect

## References

1. Deiner, K., Bik, H.M., Mächler, E., Seymour, M., Lacoursière-Roussel, A., Altermatt, F., Creer, S., Bista, I., Lodge, D.M., de Vere, N., et al. (2017). Environmental DNA metabarcoding: Transforming how we survey animal and plant communities. Mol. Ecol. 26, 5872–5895. 10.1111/mec.14350.

2. Jeunen, G.-J., Knapp, M., Spencer, H.G., Lamare, M.D., Taylor, H.R., Stat, M., Bunce, M., and Gemmell, N.J. (2019). Environmental DNA (eDNA) metabarcoding reveals strong discrimination among diverse marine habitats connected by water movement. Mol. Ecol. Resour. 19, 426–438. 10.1111/1755-0998.12982.

3. de Santana, C.D., Parenti, L.R., Dillman, C.B., Coddington, J.A., Bastos, D.A., Baldwin, C.C., Zuanon, J., Torrente-Vilara, G., Covain, R., Menezes, N.A., et al. (2021). The critical role of natural history museums in advancing eDNA for biodiversity studies: a case study with Amazonian fishes. Sci. Rep. 11, 18159. 10.1038/s41598-021-97128-3.

4. Raxworthy, C.J., and Smith, B.T. (2021). Mining museums for historical DNA: advances and challenges in museomics. Trends Ecol. Evol. 36, 1049–1060. 10.1016/j.tree.2021.07.009.

5. Gold, Z., Kelly, R.P., Shelton, A.O., Thompson, A.R., Goodwin, K.D., Gallego, R., Parsons, K.M., Thompson, L.R., Kacev, D., and Barber, P.H. (2022). Message in a Bottle: Archived DNA Reveals Marine Heatwave-Associated Shifts in Fish Assemblages (Ecology) 10.1101/2022.07.27.501788.

6. Heindler, F.M., Christiansen, H., Frédérich, B., Dettaï, A., Lepoint, G., Maes, G.E., Van de Putte, A.P., and Volckaert, F.A.M. (2018). Historical DNA Metabarcoding of the Prey and Microbiome of Trematomid Fishes Using Museum Samples. Front. Ecol. Evol. 6.

7. Webster, M.S. (2017). The Extended Specimen: Emerging Frontiers in Collections-Based Ornithological Research (CRC Press).

8. Lendemer, J., Thiers, B., Monfils, A.K., Zaspel, J., Ellwood, E.R., Bentley, A., LeVan, K., Bates, J., Jennings, D., Contreras, D., et al. (2020). The Extended Specimen Network: A Strategy to Enhance US Biodiversity Collections, Promote Research and Education. BioScience 70, 23–30. 10.1093/biosci/biz140.

9. Teixeira-Costa, L., Heberling, J.M., Wilson, C.A., and Davis, C.C. (2023). Parasitic flowering plant collections embody the extended specimen. Methods Ecol. Evol. 14, 319–331. 10.1111/2041-210X.13866.

10. Mariani, S., Baillie, C., Colosimo, G., and Riesgo, A. (2019). Sponges as natural environmental DNA samplers. Curr. Biol. 29, R401–R402. 10.1016/j.cub.2019.04.031.

11. Jeunen, G.-J., Cane, J.S., Ferreira, S., Strano, F., von Ammon, U., Cross, H., Day, R., Hesseltine, S., Ellis, K., Urban, L., et al. Assessing the utility of marine filter feeders for environmental DNA (eDNA) biodiversity monitoring. Mol. Ecol. Resour. n/a. 10.1111/1755-0998.13754.

12. Cai, W., Harper, L.R., Neave, E.F., Shum, P., Craggs, J., Arias, M.B., Riesgo, A., and Mariani, S. Environmental DNA persistence and fish detection in captive sponges. Mol. Ecol. Resour. n/a. 10.1111/1755-0998.13677.

13. Costello, M.J., Tsai, P., Wong, P.S., Cheung, A.K.L., Basher, Z., and Chaudhary, C. (2017). Marine biogeographic realms and species endemicity. Nat. Commun. 8, 1057. 10.1038/s41467-017-01121-2.

14. Taberlet, P., Bonin, A., Zinger, L., and Coissac, E. (2018). Environmental DNA: For Biodiversity Research and Monitoring (Oxford University Press).

15. Weisz, J.B., Lindquist, N., and Martens, C.S. (2008). Do associated microbial abundances impact marine demosponge pumping rates and tissue densities? Oecologia 155, 367–376. 10.1007/s00442-007-0910-0.

16. Kutti, T., Bannister, R.J., and Fosså, J.H. (2013). Community structure and ecological function of deep-water sponge grounds in the Traenadypet MPA—Northern Norwegian continental shelf. Cont. Shelf Res. 69, 21–30. 10.1016/j.csr.2013.09.011.

17. Bart, M.C., Mueller, B., Rombouts, T., van de Ven, C., Tompkins, G.J., Osinga, R., Brussaard, C.P.D., MacDonald, B., Engel, A., Rapp, H.T., et al. (2021). Dissolved organic carbon (DOC) is essential to balance the metabolic demands of four dominant North-Atlantic deep-sea sponges. Limnol. Oceanogr. 66, 925–938. 10.1002/lno.11652.

18. Lynggaard, C., Oceguera-Figueroa, A., Kvist, S., Gilbert, M.T.P., and Bohmann, K. (2022). The potential of aquatic bloodfeeding and nonbloodfeeding leeches as a tool for iDNA characterisation. Mol. Ecol. Resour. 22, 539–553. 10.1111/1755-0998.13486.

19. Siegenthaler, A., Wangensteen, O.S., Soto, A.Z., Benvenuto, C., Corrigan, L., and Mariani, S. (2019). Metabarcoding of shrimp stomach content: Harnessing a natural sampler for fish biodiversity monitoring. Mol. Ecol. Resour. 19, 206–220. 10.1111/1755-0998.12956.

20. Christiansen, S., Klevjer, T.A., Røstad, A., Aksnes, D.L., and Kaartvedt, S. (2021). Flexible behaviour in a mesopelagic fish (Maurolicus muelleri). ICES J. Mar. Sci. 78, 1623–1635. 10.1093/icesjms/fsab075.

21. Link, J.S., and Garrison, L.P. (2002). Trophic ecology of Atlantic cod Gadus morhua on the northeast US continental shelf. Mar. Ecol. Prog. Ser. 227, 109–123. 10.3354/meps227109.

22. Garcia, E.G. (2007). The Northern Shrimp (Pandalus borealis) Offshore Fishery in the Northeast Atlantic. In Advances in Marine Biology (Academic Press), pp. 147–266. 10.1016/S0065-2881(06)52002-4.

23. Albert, O.T. (1993). Distribution, population structure and diet of silvery pout (Gadiculus argenteus thori J. Schmidt), poor cod (Trisopterus minutus minutus (L.)), four-bearded rockling (Rhinonemus cimbrius (L.)), and Vahl’s eelpout (Lycodes vahlii gracilis Reinhardt) in the Norwegian Deep. Sarsia 78, 141–154. 10.1080/00364827.1993.10413531.

24. Kenchington, E., Power, D., and Koen-Alonso, M. (2013). Associations of demersal fish with sponge grounds on the continental slopes of the northwest Atlantic. Mar. Ecol. Prog. Ser. 477, 217–230. 10.3354/meps10127.

25. Sinniger, F., Pawlowski, J., Harii, S., Gooday, A.J., Yamamoto, H., Chevaldonné, P., Cedhagen, T., Carvalho, G., and Creer, S. (2016). Worldwide Analysis of Sedimentary DNA Reveals Major Gaps in Taxonomic Knowledge of Deep-Sea Benthos. Front. Mar. Sci. 3.

26. Thomsen, P.F., Møller, P.R., Sigsgaard, E.E., Knudsen, S.W., Jørgensen, O.A., and Willerslev, E. (2016). Environmental DNA from Seawater Samples Correlate with Trawl Catches of Subarctic, Deepwater Fishes. PLOS ONE 11, e0165252. 10.1371/journal.pone.0165252.

27. Everett, M.V., and Park, L.K. (2018). Exploring deep-water coral communities using environmental DNA. Deep Sea Res. Part II Top. Stud. Oceanogr. 150, 229–241. 10.1016/j.dsr2.2017.09.008.

28. Canals, O., Mendibil, I., Santos, M., Irigoien, X., and Rodríguez-Ezpeleta, N. (2021). Vertical stratification of environmental DNA in the open ocean captures ecological patterns and behavior of deep-sea fishes. Limnol. Oceanogr. Lett. 6, 339–347. 10.1002/lol2.10213.

29. Visser, F., Merten, V.J., Bayer, T., Oudejans, M.G., de Jonge, D.S.W., Puebla, O., Reusch, T.B.H., Fuss, J., and Hoving, H.J.T. (2021). Deep-sea predator niche segregation revealed by combined cetacean biologging and eDNA analysis of cephalopod prey. Sci. Adv. 7, eabf5908. 10.1126/sciadv.abf5908.

30. Fujiwara, Y., Tsuchida, S., Kawato, M., Masuda, K., Sakaguchi, S.O., Sado, T., Miya, M., and Yoshida, T. (2022). Detection of the Largest Deep-Sea-Endemic Teleost Fish at Depths of Over 2,000 m Through a Combination of eDNA Metabarcoding and Baited Camera Observations. Front. Mar. Sci. 9, 945758. 10.3389/fmars.2022.945758.

31. McClenaghan, B., Fahner, N., Cote, D., Chawarski, J., McCarthy, A., Rajabi, H., Singer, G., and Hajibabaei, M. (2020). Harnessing the power of eDNA metabarcoding for the detection of deepsea fishes. PLOS ONE 15, e0236540. 10.1371/journal.pone.0236540.

32. Taboada, S., Ríos, P., Mitchell, A., Cranston, A., Busch, K., Tonzo, V., Cárdenas, P., Sánchez, F., Leiva, C., and Koutsouveli, V. (2022). Genetic diversity, gene flow and hybridization in fan-shaped sponges (Phakellia spp.) in the North-East Atlantic deep sea. Deep Sea Res. Part Oceanogr. Res. Pap. 181, 103685.

33. Taboada, S., Whiting, C., Wang, S., Ríos, P., Davies, A., Mienis, F., Kenchington, E., Cárdenas, P., Cranston, A., and Koutsouveli, V. (2022). Connectivity of sponge grounds in the deep sea: genetic diversity, gene flow and oceanographic pathways in the fan-shaped sponge Phakellia ventilabrum in the northeast Atlantic. Authorea Prepr.

34. Pusceddu, A., Bianchelli, S., Martín, J., Puig, P., Palanques, A., Masqué, P., and Danovaro, R. (2014). Chronic and intensive bottom trawling impairs deep-sea biodiversity and ecosystem functioning. Proc. Natl. Acad. Sci. 111, 8861–8866. 10.1073/pnas.1405454111.

35. Van Dover, C.L., Ardron, J.A., Escobar, E., Gianni, M., Gjerde, K.M., Jaeckel, A., Jones, D.O.B., Levin, L.A., Niner, H.J., Pendleton, L., et al. (2017). Biodiversity loss from deep-sea mining. Nat. Geosci. 10, 464–465. 10.1038/ngeo2983.

36. Danovaro, R., Fanelli, E., Aguzzi, J., Billett, D., Carugati, L., Corinaldesi, C., Dell’Anno, A., Gjerde, K., Jamieson, A.J., Kark, S., et al. (2020). Ecological variables for developing a global deep-ocean monitoring and conservation strategy. Nat. Ecol. Evol. 4, 181–192. 10.1038/s41559-019-1091-z.

37. Maldonado, M., Aguilar, R., Bannister, R.J., Bell, J.J., Conway, K.W., Dayton, P.K., Díaz, C., Gutt, J., Kelly, M., Kenchington, E.L.R., et al. (2015). Sponge Grounds as Key Marine Habitats: A Synthetic Review of Types, Structure, Functional Roles, and Conservation Concerns. In Marine Animal Forests, S. Rossi, L. Bramanti, A. Gori, and C. Orejas Saco del Valle, eds. (Springer International Publishing), pp. 1–39. 10.1007/978-3-319-17001-5_24-1.

38. Danovaro, R., Gambi, C., Dell’Anno, A., Corinaldesi, C., Fraschetti, S., Vanreusel, A., Vincx, M., and Gooday, A.J. (2008). Exponential Decline of Deep-Sea Ecosystem Functioning Linked to Benthic Biodiversity Loss. Curr. Biol. 18, 1–8. 10.1016/j.cub.2007.11.056.

39. Harper, L.R., Neave, E.F., Sellers, G.S., Cunnington, A.V., Arias, M.B., Craggs, J., MacDonald, B., Riesgo, A., and Mariani, S. (2022). Optimised DNA isolation from marine sponges for natural sampler DNA (nsDNA) metabarcoding. 2022.07.11.499619. 10.1101/2022.07.11.499619.

40. Boyer, F., Mercier, C., Bonin, A., Le Bras, Y., Taberlet, P., and Coissac, E. (2016). obitools: a unixinspired software package for DNA metabarcoding. Mol. Ecol. Resour. 16, 176–182. 10.1111/1755-0998.12428.

41. Rognes, T., Flouri, T., Nichols, B., Quince, C., and Mahé, F. (2016). VSEARCH: a versatile open source tool for metagenomics. PeerJ 4, e2584. 10.7717/peerj.2584.

42. Mahé, F., Rognes, T., Quince, C., de Vargas, C., and Dunthorn, M. (2015). Swarm v2: highly-scalable and high-resolution amplicon clustering. PeerJ 3, e1420. 10.7717/peerj.1420.

43. Gao, X., Lin, H., Revanna, K., and Dong, Q. (2017). A Bayesian taxonomic classification method for 16S rRNA gene sequences with improved species-level accuracy. BMC Bioinformatics 18, 247. 10.1186/s12859-017-1670-4.

44. R Core Team (2022). R: A language and environment for statistical computing. R Foundation for Statistical Computing, Vienna, Austria. URL https://www.R-project.org/.

45. Jari Oksanen, F. Guillaume Blanchet, Michael Friendly, Roeland Kindt, Pierre Legendre, Dan McGlinn Peter R. Minchin, R. B. O’Hara, Gavin L. Simpson, Peter Solymos, M. Henry H. Stevens, Eduard Szoecs and Helene Wagner (2020). vegan: Community Ecology Package. R package version 2.5-7. https://CRAN.R-project.org/package=vegan

46. Benjamini, Y., and Yekutieli, D. (2001). The control of the false discovery rate in multiple testing under dependency. Annals of Statistics, 29, 1165–1188. doi:10.1214/aos/1013699998.

47. Kindt, R., and Coe, R. (2005). Tree diversity analysis: a manual and software for common statistical methods for ecological and biodiversity studies (World Agrofirestry Centre).

48. Cáceres, M.D., and Legendre, P. (2009). Associations between species and groups of sites: indices and statistical inference. Ecology 90, 3566–3574. 10.1890/08-1823.1.

49. Wickham, H., Averick, M., Bryan, J., Chang, W., McGowan, L.D., François, R., Grolemund, G., Hayes, A., Henry, L., Hester, J., et al. (2019). Welcome to the Tidyverse. J. Open Source Softw. 4, 1686. 10.21105/joss.01686.

50. H. Wickham. ggplot2: Elegant Graphics for Data Analysis. Springer-Verlag New York, 2016.

