## Supplemental for "Trapped DNA fragments in marine sponge specimens unveil north Atlantic deep-sea fish diversity"

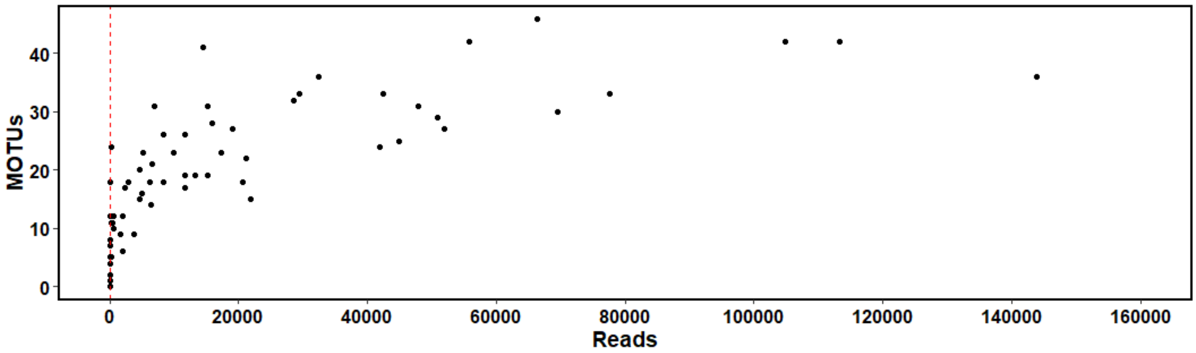


**Supplemental Figure 1.** Rarefaction curve with a dashed red line through x = 100 reads.

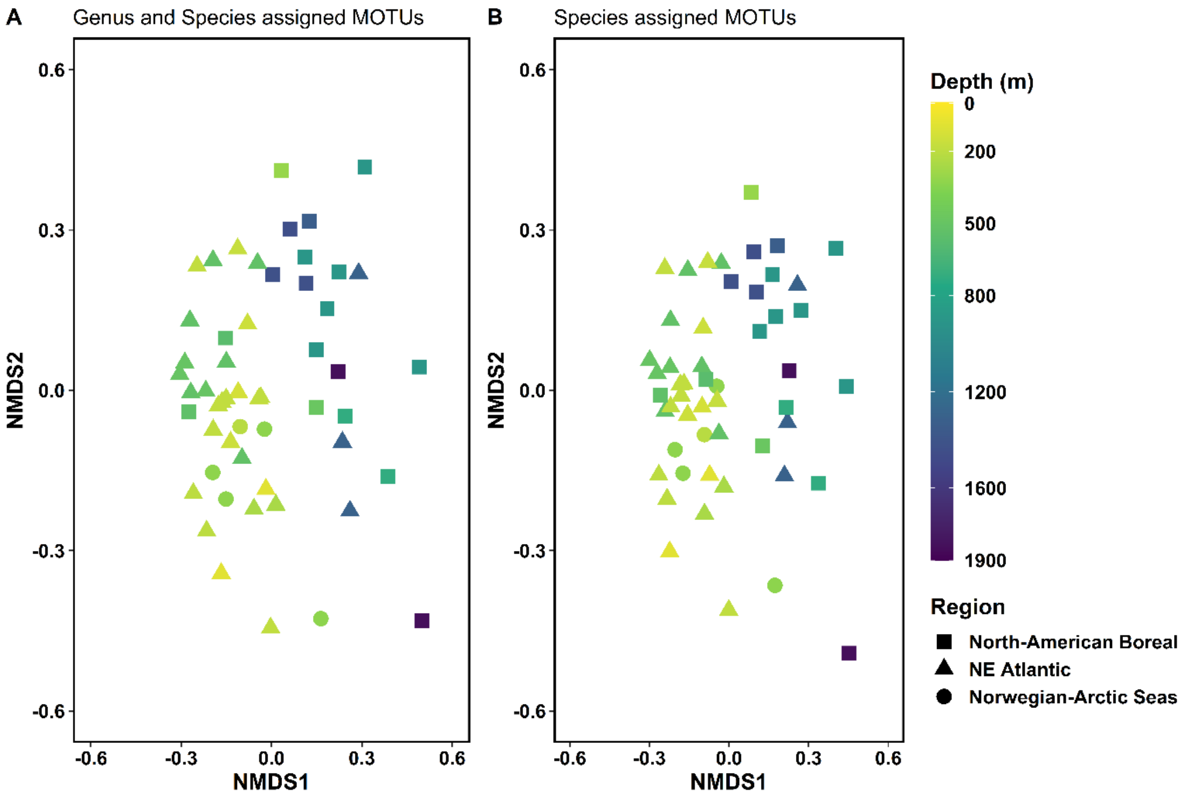


**Supplemental Figure 2.** Non-metric Multi-Dimensional Scaling (NMDS) plots conveying beta diversity from teleost and elasmobranch detections. **A** NMDS plot of a Jaccard dissimilarity matrix containing all teleost and elasmobranch MOTUs which could be assigned to either genus or species level (88). **B** NMDS plot of a Jaccard dissimilarity matrix containing all teleost and elasmobranch MOTUs which could be assigned to species level (65) (i.e., the same data plotted in Figure 2A).

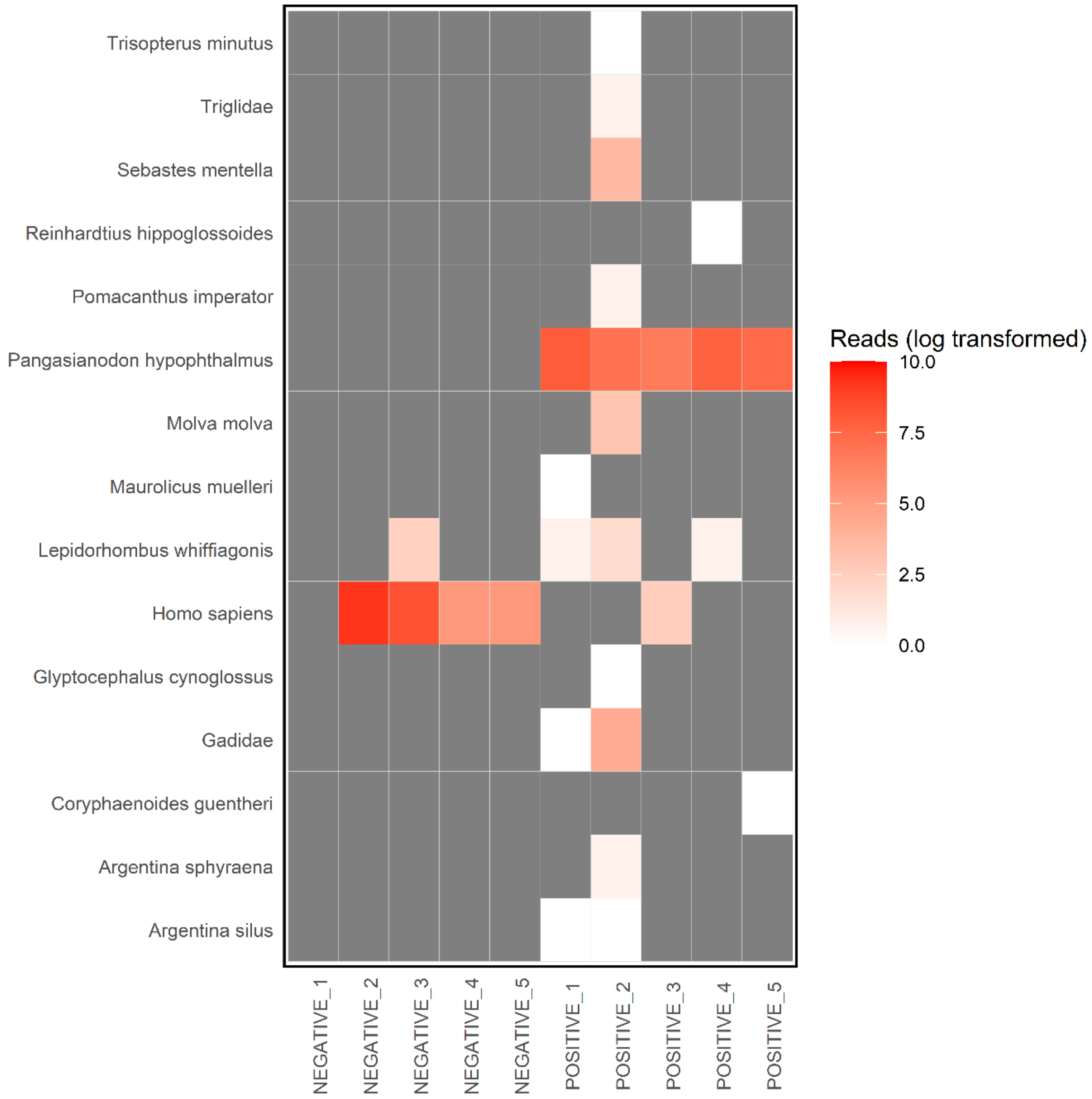


**Supplemental Figure 3.** Heat map showing log-transformed reads in controls. An explanation of how contamination was dealt with can be found in the Methods section of the main manuscript.

**Supplemental Table 1. List of sponge specimens analysed (N = 54),** where Gb = *Geodia barretti*, Gh = *Geodia henstcheli*, and Pv = *Phakellia ventilabrum.* The sponge abbreviations are used both in the 'Short ID' column and the 'Species' column. Other abbreviations: NHM = Natural History Museum (London), Lat = latitude in decimal degrees, Lon = longitude in decimal degrees, MPA = marine protected area, and UK = unknown.

| short ID | long ID | NHM Voucher # (in progress) | Species | Aggregation | Region | Lat | Lon | Depth (m) | MPA Status | Month | Year |
|---|---|---|---|---|---|---|---|---|---|---|---|
| 1.Gb | Gb_FS_MPA_14N4_s_50 | | Gb | Faroe Shetland Sponge Belt | NE Atlantic | 60.83583 | -3.03717 | 525 | MPA | August | 2018 |
| 2.Gb | Gb_FS_MPA_20T6_s_49 | | Gb | Faroe Shetland Sponge Belt | NE Atlantic | 60.83583 | -3.03717 | 525 | MPA | August | 2018 |
| 3.Gb | Gb_FS_MPA_22V2_s_51 | | Gb | Faroe Shetland Sponge Belt | NE Atlantic | 60.83583 | -3.03717 | 525 | MPA | August | 2018 |
| 4.Gb | Gb_RSB_MPA_13M4_d_74 | | Gb | W Rosemary bank | NE Atlantic | 59.12883 | -10.8078 | 1304 | MPA | September | 2016 |
| 5.Gb | Gb_RSB_MPA_13M5_d_72 | | Gb | W Rosemary bank | NE Atlantic | 59.12883 | -10.8078 | 1304 | MPA | September | 2016 |
| 6.Gb | Gb_RSB_MPA_18R4_d_73 | | Gb | W Rosemary bank | NE Atlantic | 59.12883 | -10.8078 | 1304 | MPA | September | 2016 |
| 7.Gb | Gb_Sva_20T5_s_25 | | Gb | Trondheimsfjorden | NE Atlantic | 63.58612 | 9.84555 | 191 | No MPA | UK | 2016 |
| 8.Pv | Pv_FS_MPA_01601_s_52 | | Pv | Faroe Shetland Sponge Belt | NE Atlantic | 60.83583 | -3.03717 | 525 | MPA | August | 2018 |
| 9.Pv | Pv_FS_MPA_01602_s_26 | | Pv | Faroe Shetland Sponge Belt | NE Atlantic | 60.83583 | -3.03717 | 525 | MPA | August | 2018 |
| 10.Pv | Pv_FS_MPA_01603_s_53 | | Pv | Faroe Shetland Sponge Belt | NE Atlantic | 60.83583 | -3.03717 | 525 | MPA | August | 2018 |
| 11.Pv | Pv_NoS_11452_s_08 | | Pv | North of Shetland | NE Atlantic | 60.454 | 0.538 | 131 | No MPA | January | 2019 |
| 12.Pv | Pv_NoS_11456_s_22 | | Pv | North of Shetland | NE Atlantic | 61.116 | 0.175 | 155 | No MPA | January | 2019 |
| 13.Pv | Pv_NoS_Notag_s_09 | | Pv | North of Shetland | NE Atlantic | 61.116 | 0.175 | 155 | No MPA | January | 2019 |
| 14.Pv | Pv_NwK_ST5_38_s_29 | | Pv | Norway | NE Atlantic | 59.81359 | 5.603875 | 259 | No MPA | September | 2016 |
| 15.Pv | Pv_NwK_ST5_39_s_30 | | Pv | Norway | NE Atlantic | 59.81359 | 5.603875 | 259 | No MPA | September | 2016 |

| | | | | | | | | | | |
|---|---|---|---|---|---|---|---|---|---|---|
| 16.Pv | Pv_RB_11218_s_13 | Pv | Rockall Bank | NE Atlantic | 57.52475 | -13.4195 | 173.5 | No MPA | September | 2018 |
| 17.Pv | Pv_RB_11221_s_23 | Pv | Rockall Bank | NE Atlantic | 57.52475 | -13.4195 | 173.5 | No MPA | September | 2018 |
| 18.Pv | Pv_RB_1418_s_15 | Pv | Rockall Bank | NE Atlantic | 57.1705 | -13.638 | 187 | No MPA | May | 2018 |
| 19.Pv | Pv_RB_1420_s_16 | Pv | Rockall Bank | NE Atlantic | 57.1705 | -13.638 | 187 | No MPA | May | 2018 |
| 20.Pv | Pv_RB_MPA_11417_s_20 | Pv | Rockall Bank | NE Atlantic | 56.7755 | -14.4681 | 189.5 | MPA | September | 2018 |
| 21.Pv | Pv_RB_MPA_11419_s_21 | Pv | Rockall Bank | NE Atlantic | 56.7755 | -14.4681 | 189.5 | MPA | September | 2018 |
| 22.Pv | Pv_RB_MPA_1865_s_24 | Pv | Rockall Bank | NE Atlantic | 56.905 | -14.741 | 186 | MPA | April | 2018 |
| 23.Pv | Pv_RB_MPA_1866_s_14 | Pv | Rockall Bank | NE Atlantic | 56.905 | -14.741 | 186 | MPA | April | 2018 |
| 24.Pv | Pv_SRT_046_26_s_11 | Pv | Sula Reef | NE Atlantic | 63.65222 | 9.758833 | 203 | No MPA | October | 2016 |
| 25.Pv | Pv_SRT_046_49_s_12 | Pv | Sula Reef | NE Atlantic | 63.65222 | 9.758833 | 203 | No MPA | October | 2016 |
| 26.Pv | Pv_SSNd_H044_01642_s_45 | Pv | Shetland Shelf North | NE Atlantic | 60.106 | -4.808 | 496 | No MPA | May | 2018 |
| 27.Pv | Pv_SSNd_H044_01643_s_46 | Pv | Shetland Shelf North | NE Atlantic | 60.106 | -4.808 | 496 | No MPA | May | 2018 |
| 28.Pv | Pv_SSNd_H044_01645_s_47 | Pv | Shetland Shelf North | NE Atlantic | 60.106 | -4.808 | 496 | No MPA | May | 2018 |
| 29.Pv | Pv_Sw_11_s_05 | Pv | Sweden | NE Atlantic | 58.89083 | 11.09502 | 90 | No MPA | March | 2019 |
| 30.Pv | Pv_Sw_9_s_04 | Pv | Sweden | NE Atlantic | 58.89083 | 11.09502 | 90 | No MPA | March | 2019 |
| 31.Gb | Gb_Can_M_Trip8_75.1_d_69 | Gb | Arctic NAFO 0A_0B (close to Paamiut) | North American Boreal | 66.80795 | -58.6674 | 903 | No MPA | October | 2016 |
| 32.Gb | Gb_Can_M_Trip8_75.2_d_70 | Gb | Arctic NAFO 0A_0B (close to Paamiut) | North American Boreal | 66.80795 | -58.6674 | 903 | No MPA | October | 2016 |
| 33.Gb | Gb_Can_M_Trip8_75.3_d_71 | Gb | Arctic NAFO 0A_0B (close to Paamiut) | North American Boreal | 66.80795 | -58.6674 | 903 | No MPA | October | 2016 |

| | | | | | | | | | | |
|---|---|---|---|---|---|---|---|---|---|---|
| 34.Gb | Gb_Can_S_15O6_d_75 | Gb | Davis Strait | North American Boreal | 62.5184 | -59.9705 | 1332 | No MPA | October | 2014 |
| 35.Gb | Gb_Can_S_16P6_s_55 | Gb | Davis Strait | North American Boreal | 62.56012 | -61.3943 | 568 | No MPA | October | 2014 |
| 36.Gb | Gb_Can_S_17Q3_d_77 | Gb | Davis Strait | North American Boreal | 62.0539 | -60.0918 | 1427 | No MPA | October | 2014 |
| 37.Gb | Gb_Can_S_23W6_d_78 | Gb | Davis Strait | North American Boreal | 61.8989 | -60.1364 | 1440 | No MPA | October | 2014 |
| 38.Gb | Gb_Can_S_2B6_d_76 | Gb | Davis Strait | North American Boreal | 62.0539 | -60.0918 | 1427 | No MPA | October | 2014 |
| 39.Gb | Gb_Can_S_5E2_s_44 | Gb | Davis Strait | North American Boreal | 62.19923 | -61.2192 | 483 | No MPA | October | 2014 |
| 40.Gb | Gb_Can_S_8H6_s_54 | Gb | Davis Strait | North American Boreal | 63.26382 | -60.3552 | 536 | No MPA | October | 2014 |
| 41.Gh | Gh_Can_M_Trip8_75.5_d_66 | Gh | Arctic NAFO 0A_0B (close to Paamiut) | North American Boreal | 66.80795 | -58.6674 | 903 | No MPA | October | 2016 |
| 42.Gh | Gh_Can_M_Trip8_75.7_d_67 | Gh | Arctic NAFO 0A_0B (close to Paamiut) | North American Boreal | 66.80795 | -58.6674 | 903 | No MPA | October | 2016 |
| 43.Gh | Gh_Can_M_Trip8_75.8_d_68 | Gh | Arctic NAFO 0A_0B (close to Paamiut) | North American Boreal | 66.80795 | -58.6674 | 903 | No MPA | October | 2016 |
| 44.Gh | Gh_Ice_A11_641_B_s_25 | Gh | Iceland | North American Boreal | 66.63883 | -12.7558 | 308.5 | No MPA | UK | 2019 |
| 45.Gh | Gh_Ice_ME85_3_1123_2_C_d_59 | Gh | Iceland | North American Boreal | 67.214 | -26.1992 | 723.35 | No MPA | September | 2011 |

| | | | | | | | | | | |
|---|---|---|---|---|---|---|---|---|---|---|
| 46.Gh | Gh_Ice_ME85_3_1123_B_d_60 | Gh | Iceland | North American Boreal | 67.214 | -26.1992 | 723.35 | No MPA | September | 2011 |
| 47.Gh | Gh_JMR_SM_AGT03_26_d_93 | Gh | Between Jan Mayen_Schultz Massif (Mohn) | North American Boreal | 71.30333 | -5.28733 | 1848.5 | No MPA | July | 2014 |
| 48.Gh | Gh_JMR_SM_AGT03_27_d_94 | Gh | Between Jan Mayen_Schultz Massif (Mohn) | North American Boreal | 71.30333 | -5.28733 | 1848.5 | No MPA | July | 2014 |
| 49.Gh | Gh_Vst_019TVG_010B_d_91 | Gh | Vesterisbanken | North American Boreal | 73.61406 | -9.03518 | 1784 | No MPA | August | 2019 |
| 50.Gb | Gb_BS_imp_9I4_s_39 | Gb | Barents Sea | Norwegian and Arctic Seas | 71.58797 | 21.37533 | 333 | No MPA | March | 2017 |
| 51.Gb | Gb_Sva_7G6_s_26 | Gb | Svalbard | Norwegian and Arctic Seas | 80.53853 | 15.36161 | 215 | No MPA | September | 2011 |
| 52.Pv | Pv_SMK_ROV25_12_s_33 | Pv | Tromsø shelf | Norwegian and Arctic Seas | 71.39517 | 16.81599 | 330 | No MPA | UK | 2017 |
| 53.Pv | Pv_SMK_ROV25_13_s_35 | Pv | Tromsø shelf | Norwegian and Arctic Seas | 71.39517 | 16.81599 | 330 | No MPA | UK | 2017 |
| 54.Pv | Pv_SMK_ROV25_14_s_36 | Pv | Tromsø shelf | Norwegian and Arctic Seas | 71.39517 | 16.81599 | 330 | No MPA | UK | 2017 |

**Supplemental Table 2. Reads from (N = 74) samples sequenced, including controls (N = 10) and samples which were removed from the statistical analysis for having low reads (N = 10).**

| | Reads |
|---|---|
| **Total** | 5,269,740 |
| Total after 'illuminapairedend' step with Q40 threshold | 4,565,067 |
| Total after bioinformatic pipeline | 3,091,641 |
| Total assigned taxonomy (Eukaryota) | 3,040,133 |
| **Total assigned to target taxa (Actinopterygii)** | 2,531,138 |
| Total assigned to species-level (99% simlaity) Actinopterygii | 1,778,145 |
| Total assigned to species-level (99% simlaity) Actinopterygii, Elasmobranchii, Chondrichthyes | 1,781,183 |
| Total assigned to species-level (99% simlaity) Actinopterygii, Elasmobranchii, Chondrichthyes, Mammalia marine-only | 1,782,119 |

**Supplemental Table 3. Fish identified to species level at a similarity threshold of ≥ 99%, alphabetically by class**.

| Class | Order | Family | Common name | Scientific name | No. of Samples Detected |
|---|---|---|---|---|---|
| Actinopteri | Anguilliformes | Nettastomatidae | Blackfin sorcerer | *Nettastoma melanurum* | 1 |
| Actinopteri | Anguilliformes | Synaphobranchidae | Kaup's arrow-tooth eel | *Synaphobranchus kaupii* | 6 |
| Actinopteri | Argentiniformes | Argentinidae | Greater argentine | *Argentina silus* | 46 |
| Actinopteri | Argentiniformes | Argentinidae | Argentine | *Argentina sphyraena* | 34 |
| Actinopteri | Argentiniformes | Bathylagidae | Slender blacksmelt | *Bathylagus pacificus* | 10 |
| Actinopteri | Aulopiformes | Bathysauridae | Deepsea lizardfish | *Bathysaurus ferox* | 1 |
| Actinopteri | Aulopiformes | Chlorophthalmidae | Shortnose greeneye | *Chlorophthalmus agassizi* | 5 |
| Actinopteri | Aulopiformes | Paralepididae | spotted barracudina | *Arctozenus risso* | 3 |
| Actinopteri | Beloniformes | Belonidae | Garfish | *Belone belone* | 1 |
| Actinopteri | Beloniformes | Exocoetidae | Tropical two-wing flyingfish | *Exocoetus volitans* | 3 |
| Actinopteri | Caproiformes | Caproidae | Boarfish | *Capros aper* | 5 |
| Actinopteri | Carangiformes | Carangidae | Atlantic horse mackerel | *Trachurus trachurus* | 17 |
| Actinopteri | Clupeiformes | Clupeidae | Atlantic herring | *Clupea harengus* | 19 |
| Actinopteri | Clupeiformes | Clupeidae | European pilchard | *Sardina pilchardus* | 3 |
| Actinopteri | Clupeiformes | Clupeidae | European sprat | *Sprattus sprattus* | 4 |
| Actinopteri | Gadiformes | Gadidae | Norway pout | *Trisopterus esmarkii* | 31 |
| Actinopteri | Gadiformes | Gadidae | Poor cod | *Trisopterus minutus* | 44 |
| Actinopteri | Gadiformes | Gadidae | Blue whiting | *Micromesistius poutassou* | 33 |
| Actinopteri | Gadiformes | Gadidae | Pouting | *Trisopterus luscus* | 1 |
| Actinopteri | Gadiformes | Gadidae | Atlantic cod | *Gadus morhua* | 33 |
| Actinopteri | Gadiformes | Gadidae | Pollock; Saithe | *Pollachius virens* | 22 |
| Actinopteri | Gadiformes | Gadidae | Silvery pout | *Gadiculus argenteus* | 27 |
| Actinopteri | Gadiformes | Lotidae | Blue ling | *Molva dypterygia* | 14 |

| | | | | | |
|---|---|---|---|---|---|
| Actinopteri | Gadiformes | Lotidae | Common ling | *Molva molva* | 36 |
| Actinopteri | Gadiformes | Lotidae | Cusk | *Brosme brosme* | 10 |
| Actinopteri | Gadiformes | Macrouridae | Günther's grenadier | *Coryphaenoides guentheri* | 10 |
| Actinopteri | Gadiformes | Macrouridae | Mediterranean grenadier | *Coryphaenoides mediterraneus* | 1 |
| Actinopteri | Gadiformes | Macrouridae | Roundnose grenadier | *Coryphaenoides rupestris* | 10 |
| Actinopteri | Gadiformes | Macrouridae | Shortbeard grenadier | *Coryphaenoides brevibarbis* | 1 |
| Actinopteri | Gadiformes | Macrouridae | Roughhead grenadier | *Macrourus berglax* | 15 |
| Actinopteri | Gadiformes | Merlucciidae | European hake | *Merluccius merluccius* | 34 |
| Actinopteri | Gadiformes | Moridae | Slender codling | *Halargyreus johnsonii* | 2 |
| Actinopteri | Labriformes | Labridae | Cuckoo wrasse | *Labrus mixtus* | 2 |
| Actinopteri | Lophiiformes | Lophiidae | Angler | *Lophius piscatorius* | 32 |
| Actinopteri | Myctophiformes | Myctophidae | Lanternfish | *Notoscopelus elongatus* | 9 |
| Actinopteri | Myctophiformes | Myctophidae | Arctic telescope | *Protomyctophum arcticum* | 9 |
| Actinopteri | Myctophiformes | Myctophidae | Rakery beaconlamp | *Lampanyctus macdonaldi* | 5 |
| Actinopteri | Notacanthiformes | Notacanthidae | Smallmouth spiny eel | *Polyacanthonotus rissoanus* | 4 |
| Actinopteri | Perciformes | Cyclopteridae | Lumpfish | *Cyclopterus lumpus* | 1 |
| Actinopteri | Perciformes | Gasterosteidae | Three-spined stickleback | *Gasterosteus aculeatus* | 1 |
| Actinopteri | Perciformes | Liparidae | Threadfin seasnail | *Rhodichthys regina* | 2 |
| Actinopteri | Perciformes | Sebastidae | Beaked redfish | *Sebastes mentella* | 51 |
| Actinopteri | Perciformes | Zoarcidae | Atlantic eelpout | *Lycodes terraenovae* | 17 |
| Actinopteri | Perciformes | Zoarcidae | Moray wolf eel | *Lycenchelys muraena* | 9 |
| Actinopteri | Perciformes | Zoarcidae | Vahl's eelpout | *Lycodes vahlii* | 4 |
| Actinopteri | Pleuronectiformes | Bothidae | Mediterranean scaldfish | *Arnoglossus laterna* | 2 |
| Actinopteri | Pleuronectiformes | Pleuronectidae | Witch flounder | *Glyptocephalus cynoglossus* | 18 |
| Actinopteri | Pleuronectiformes | Pleuronectidae | Greeenland halibut | *Reinhardtius hippoglossoides* | 52 |
| Actinopteri | Pleuronectiformes | Scophthalmidae | Turbot | *Scophthalmus maximus* | 6 |
| Actinopteri | Pleuronectiformes | Scophthalmidae | Megrim | *Lepidorhombus whiffiagonis* | 50 |
| Actinopteri | Pleuronectiformes | Soleidae | Common sole | *Solea solea* | 2 |
| Actinopteri | Pleuronectiformes | Soleidae | Thickback sole | *Microchirus variegatus* | 8 |
| Actinopteri | Scombriformes | Scombridae | Atlantic mackerel | *Scomber scombrus* | 48 |
| Actinopteri | Scombriformes | Trichiuridae | Black scabbardfish | *Aphanopus carbo* | 3 |
| Actinopteri | Scombriformes | Trichiuridae | Silver scabbardfish | *Lepidopus caudatus* | 3 |
| Actinopteri | Squaliformes | Etmopteridae | Black dogfish | *Centroscyllium fabricii* | 2 |
| Actinopteri | Stomiiformes | Sternoptychidae | Silvery lightfish | *Maurolicus muelleri* | 25 |
| Actinopteri | Syngnathiformes | Mullidae | Striped red mulllet | *Mullus surmuletus* | 1 |
| Actinopteri | Zeiformes | Zeidae | John Dory | *Zeus faber* | 2 |

| | | | | | |
|---|---|---|---|---|---|
| Chondrichthyes | Chimaeriformes | Chimaeridae | Rabbit fish | *Chimaera monstrosa* | 17 |
| Chondrichthyes | Chimaeriformes | Chimaeridae | Small-eyed rabbitfish fish | *Hydrolagus affinis* | 4 |
| Chondrichthyes | Chimaeriformes | Chimaeridae | Large-eyed rabbitfish | *Hydrolagus mirabilis* | 1 |
| Chondrichthyes | Carcharhiniformes | Scyliorhinidae | Lesser spotted dogfish | *Scyliorhinus canicula* | 1 |
| Chondrichthyes | Rajiformes | Rajidae | Blue skate | *Dipturus batis* | 3 |
| Chondrichthyes | Rajiformes | Rajidae | Shagreen ray | *Leucoraja fullonica* | 1 |

**Supplemental Table 4. Results of the multivariate homogeneity of group dispersions (*betadisper* and *anova* functions), permutational multivariate analysis of variance (i.e., PERMANOVA, *adonis* function), including pairwise comparisons and fits of environmental vectors (*envfit* function) of fish OTUs identified to the species level:** a-b) between groups; c-d) between groups (i.e., biogeographic region) and treatments (pairwise comparisons); e) goodness of fit of environmental vectors: latitude, depth and sampling year. Tests are based on Jaccard's dissimilarity distances and 1000 permutations. P(BH): P value corrected with the Benjamini-Hochberg method. Significant P values are bolded. All functions are available in the R package vegan.

a) Multivariate homogeneity of group dispersions analysis between biogeographic regions.

| Source | DF | Sq | Mean Sq | F Value | P value |
|---|---|---|---|---|---|
| Groups | 2 | 0.04797 | 0.023987 | 1.9433 | 0.1537 |
| Residuals | 51 | 0.62953 | 0.012344 | | |

b) Multivariate homogeneity of group dispersions analysis between sponge species.

| Source | DF | Sq | Mean Sq | F Value | P value |
|---|---|---|---|---|---|
| Groups | 2 | 0.08152 | 0.040759 | 4.8151 | **0.01215** |
| Residuals | 51 | 0.43170 | 0.008465 | | |

c) PERMANOVA between biogeographic regions.

| Source | DF | Sums of Sqs | Mean Sqs | F Model | $R^2$ | P value |
|---|---|---|---|---|---|---|
| Groups | 2 | 1.7391 | 0.86955 | 5.0062 | 0.16411 | **0.000999** |
| Residuals | 51 | 8.8584 | 0.17369 | 0.83589 | | |
| Total | 53 | 10.5975 | 1.00000 | | | |

d) PERMANOVA for pairwise comparisons between each combination of biogeographic regions.

| Source | DF | Sums of Sqs | Mean Sqs | F Model | $R^2$ | P value | P(BH) |
|---|---|---|---|---|---|---|---|
| Northeast Atlantic; North American Boreal | 1 | 1.3782 | 1.37824 | 7.8209 | 0.14266 | **0.000999** | **0.001498501** |
| Residuals | 47 | 8.2826 | 0.17622 | 0.85734 | | | |
| Total | 48 | 9.6608 | 1.00000 | | | | |
| Northeast Atlantic; Norwegian and Arctic Seas | 1 | 0.3034 | 0.30341 | 1.9246 | 0.05511 | **0.02498** | **0.024975025** |
| Residuals | 33 | 5.2025 | 0.15765 | 0.94489 | | | |
| Total | 34 | 5.5059 | 1.00000 | | | | |
| North American Boreal; Norwegian and Arctic Seas | 1 | 0.6566 | 0.65659 | 3.4135 | 0.13432 | **0.000999** | **0.001498501** |
| Residuals | 22 | 4.2317 | 0.19235 | 0.86568 | | | |
| Total | 23 | 4.8883 | 1.00000 | | | | |

e) Correlation of environmental vectors to site ordination scores.

| Vector | NMDS1 | NMDS2 | $R^2$ | P value |
|---|---|---|---|---|
| Latitude | 0.91612 | -0.40091 | 0.3496 | **0.000999** |
| Sampling Depth | 0.95229 | 0.30518 | 0.5756 | **0.000999** |
| Sampling Year | -0.99819 | 0.06007 | 0.1679 | **0.017982** |

**Supplemental Table 5. Results of the multivariate homogeneity of group dispersions (*betadisper* and *anova* functions), permutational multivariate analysis of variance (i.e., PERMANOVA, *adonis* function), including pairwise comparisons:** a-b) between groups; c-d) between groups (i.e., populations, marine protected area (MPA) status), e) pairwise comparisons between populations. Tests are based on Jaccard's dissimilarity distances and 1000 permutations. P(BH): P value corrected with the Benjamini-Hochberg method. Significant P values are bolded. All functions are available in the R package vegan.

a) Multivariate homogeneity of group dispersions analysis between *P. ventilabrum* populations.

| Source | DF | Sq | Mean Sq | F Value | P value |
|---|---|---|---|---|---|
| Groups | 6 | 0.058586 | 0.0097643 | 0.927 | 0.5018 |
| Residuals | 16 | 0.168533 | 0.0105333 | | |

b) Multivariate homogeneity of group dispersions analysis between *P. ventilabrum* specimens collected within verses outside MPAs.

| Source | DF | Sq | Mean Sq | F Value | P value |
|---|---|---|---|---|---|
| Groups | 1 | 0.003008 | 0.0030083 | 0.4077 | 0.5301 |
| Residuals | 21 | 0.154968 | 0.0073795 | | |

c) PERMANOVA between *P. ventilabrum* populations.

| Source | DF | Sums of Sqs | Mean Sqs | F Model | $R^2$ | P value |
|---|---|---|---|---|---|---|
| Groups | 6 | 1.5167 | 0.252779 | 2.7152 | 0.50451 | **0.000999** |
| Residuals | 16 | 8.8584 | 0.17369 | 0.83589 | | |
| Total | 22 | 10.5975 | 1.00000 | | | |

d) PERMANOVA between *P. ventilabrum* specimens collected within verses outside MPAs.

| Source | DF | Sums of Sqs | Mean Sqs | F Model | $R^2$ | P value |
|---|---|---|---|---|---|---|
| Groups | 1 | 0.26186 | 0.26186 | 2.0038 | 0.08711 | **0.02597** |
| Residuals | 21 | 2.74436 | 0.13068 | 0.91289 | | |
| Total | 22 | 3.00622 | 1.00000 | | | |

e) PERMANOVA for pairwise comparisons between each combination of *P. ventilabrum* locations. The abbreviations refer to the following sponge locations: 'fs'= Faroe Shetland Sponge Belt; 's'= North of Shetland; 'nk'= Norway; 'rb'= Rockall Bank; 'ss'= Shetland Shelf; 'tf'= Sula Reef; and 'sw'= Sweden.

| Pairwise Comparison | P value | P(BH) | Pairwise Comparison | P value | P(BH) |
|---|---|---|---|---|---|
| fs_s | 0.100000000 | 0.1400000 | nk_rb | 0.050949051 | 0.1400000 |
| fs_nk | 0.100000000 | 0.1400000 | nk_ss | 0.100000000 | 0.1400000 |
| fs_rb | **0.005994006** | 0.1258741 | nk_tf | 1.000000000 | 1.0000000 |
| fs_ss | 0.100000000 | 0.1400000 | nk_sw | 0.666666667 | 0.7000000 |
| fs_tf | 0.100000000 | 0.1400000 | rb_ss | 0.071928072 | 0.1400000 |
| fs_sw | 0.100000000 | 0.1400000 | rb_tf | 0.050949051 | 0.1400000 |
| nk_s | 0.500000000 | 0.5526316 | rb_sw | 0.053946054 | 0.1400000 |
| rb_s | 0.155844156 | 0.2045455 | ss_sw | 0.100000000 | 0.1400000 |
| ss_s | 0.100000000 | 0.1400000 | ss_tf | 0.100000000 | 0.1400000 |
| tf_s | 0.300000000 | 0.3705882 | sw_tf | 0.333333333 | 0.3888889 |
| sw_s | 0.100000000 | 0.1400000 | | | |

**Supplemental Table 6. Results of the indicator value species analysis (*multipatt* function) and multilevel pattern analysis (*IndVal.g* method):** a-b) sampling depth ranges; c-d) biogeographic regions; e-f) MPA status in the Northeast Atlantic with *P. ventilabrum* samples. Tests underwent 10000 permutations and the number of indicator species, indicator species for both depth range and biogeographic region as well as significant P values are bolded. "A" is the estimate probability that samples are associated to the matched group(s) if the indicator species has been detected in the sample (i.e., specificity or predictive value). "B" is the estimate probability of detecting the indicator species in the matched group(s) (i.e., sensitivity). "Stat" is the indicator value index which suggests the strength of the indicator species association to the group(s). Associations can be positive or negative. Groups separated with a "+" denote an additional group association to the indicator species. All functions are available in the R package Multipatt.

a) Summary table of indicator value and multilevel pattern analysis for sampling depth ranges.

| | |
|---|---|
| Total number of Species | 65 |
| Selected number of Species | **16** |
| Number of species associated to 1 group | 2 |
| " 2 groups | 5 |
| " 3 groups | 5 |
| " 4 groups | 3 |
| " 5 groups | 1 |

b) Results of the indicator value and multilevel pattern analysis for sampling depth ranges.

| Species | Depth Range Group(s) separated by '+' | A | B | Stat | P value |
|---|---|---|---|---|---|
| *Lycodes vahlii* | 500-800 | 1.0 | 0.4 | 0.632 | **0.0181** |
| *Hydrolagus affinis* | 1200-1600 | 0.8571 | 0.4286 | 0.606 | **0.208** |
| ***Bathylagus euroyops*** | 800-1200 + 1200-1600 | 0.8947 | 0.6154 | 0.742 | **0.0038** |
| *Coryphaenoides guentheri* | 800-1200 + 1200-1600 | 0.7471 | 0.4615 | 0.587 | **0.0401** |
| *Lampanyctus macdonaldi* | 800-1200 + 1200-1600 | 1.0000 | 0.3846 | 0.620 | **0.0247** |
| ***Macrourus berglax*** | 800-1200 + 1200-1600 | 0.7888 | 0.7692 | 0.779 | **0.0001** |
| *Protomyctophum arcticum* | 500-800 + 800-1200 | 0.8673 | 0.5000 | 0.659 | **0.0068** |
| *Lycodes terraenovae* | 800-1200 + 1200-1600 + >1600 | 0.7453 | 0.6250 | 0.683 | **0.0133** |
| *Clupea harengus* | 80-200 + 200-500 + 500-800 | 1.0000 | 0.5000 | 0.707 | **0.0082** |
| ***Glyptocephalus cynoglossus*** | 80-200 + 200-500 + 500-800 | 1.0000 | 0.4737 | 0.688 | **0.0135** |
| ***Pollachius virens*** | 80-200 + 200-500 + 500-800 | 1.0000 | 0.5789 | 0.761 | **0.0015** |
| ***Trisopterus esmarkii*** | 80-200 + 200-500 + 500-800 | 1.0000 | 0.8158 | 0.903 | **0.0001** |
| *Argentina sphyraena* | 80-200 + 200-500 + 500-800 + 1200-1600 | 0.9451 | 0.7333 | 0.833 | **0.0091** |
| ***Gadiculus argenteus*** | 80-200 + 200-500 + 500-800 + 1200-1600 | 1.0000 | 0.6000 | 0.775 | **0.0073** |
| ***Gadus morhua*** | 80-200 + 200-500 + 500-800 + 1200-1600 | 1.0000 | 0.7333 | 0.856 | **0.0003** |
| *Maurolicus muelleri* | 80-200 + 500-800 + 800-1200 + 1200-1600 + >1600 | 0.9489 | 0.5750 | 0.739 | **0.0274** |

c) Summary table of indicator value and multilevel pattern analysis for biogeographic regions.

| | |
|---|---|
| Total number of Species | 65 |
| Selected number of Species | **8** |
| Number of species associated to 1 group | 4 |
| " 2 groups | 4 |

d) Results of the indicator value and multilevel pattern analysis for biogeographic regions.

| Species | Group(s) separated by '+' | A | B | Stat | P value |
|---|---|---|---|---|---|
| ***Glyptocephalus cynoglossus*** | Northeast Atlantic | 0.8352 | 0.5333 | 0.667 | **0.0437** |
| Microchirus variegatus | Northeast Atlantic | 1.0000 | 0.2667 | 0.516 | **0.0381** |

| | | | | | |
|---|---|---|---|---|---|
| ***Bathylagus euroyops*** | North American Boreal | 0.8633 | 0.4211 | 0.603 | **0.0414** |
| ***Macrourus berglax*** | North American Boreal | 0.8633 | 0.6316 | 0.738 | **0.0012** |
| ***Gadiculus argenteus*** | Northeast Atlantic + Norwegian and Arctic Seas | 1.0000 | 0.7714 | 0.878 | **0.0001** |
| ***Gadus morhua*** | Northeast Atlantic + Norwegian and Arctic Seas | 0.8588 | 0.8000 | 0.829 | **0.0007** |
| ***Pollachius virens*** | Northeast Atlantic + Norwegian and Arctic Seas | 0.9629 | 0.6000 | 0.760 | **0.0092** |
| ***Trisopterus esmarkii*** | Northeast Atlantic + Norwegian and Arctic Seas | 0.9180 | 0.8000 | 0.857 | **0.0001** |

e) Summary table of indicator value and multilevel pattern analysis for MPA status in the Northeast Atlantic with *P. ventilabrum* samples.

| | |
|---|---|
| Total number of Species | 58 |
| Selected number of Species | **4** |
| Number of species associated to 1 group | 4 |

f) Results of the indicator value and multilevel pattern analysis for MPA status.

| Species | Group | A | B | Stat | P value |
|---|---|---|---|---|---|
| *Lycenchelys muraena* | MPA | 0.9999 | 0.5714 | 0.756 | **0.0069** |
| *Lycodes terraenovae* | MPA | 0.9998 | 0.5714 | 0.756 | **0.0074** |
| *Protomyctophum arcticum* | MPA | 1.0000 | 0.4286 | 0.655 | **0.0176** |
| *Lycodes vahlii* | MPA | 1.0000 | 0.4286 | 0.655 | **0.0176** |
